# 12 weeks of voluntary wheel running restores glucagon sensitivity in middle-aged mice

**DOI:** 10.1101/2025.05.23.655872

**Authors:** Tyler J. Marx, Kassandra Bruner, Susma Ghimire, Temara Fletcher, Jennifer H. Stern

**Affiliations:** College of Medicine, Department of Medicine, Division of Endocrinology, University of Arizona, Tucson, AZ, USA; Bio 5 Institute, University of Arizona, Tucson, AZ, USA

**Keywords:** Glucagon, exercise training, aging, glycogen

## Abstract

Aerobic exercise training is a potent intervention for the treatment and prevention of age-related metabolic disease, which is characterized by both insulin and glucagon resistance. While insulin resistance is a key driver of metabolic disease in aging, glucagon signaling is equally critical in maintaining both glucose and lipid homeostasis, particularly during exercise. Previous studies have established the glucagon sensitizing effects of exercise training in younger animals. Most studies in rodents employ rigorous and carefully dosed forced exercise protocols. This forced exercise is a stressful paradigm. We implemented a voluntary wheel running (VWR) intervention to assess the effects of aging and exercise training on glucagon sensitivity. We initiated 12-weeks of VWR in young adult (6-month-old) and middle-aged (12-month-old) C57BL/6NCrl male mice. Glucagon sensitivity, as assessed by glucagon stimulated hyperglycemia, was decreased in middle-aged compared to young adult sedentary mice (P=0.046). While VWR did not affect glucose clearance, circulating insulin, glucagon, or insulin sensitivity, regardless of age, VWR improved glucagon responsivity only in middle-aged mice (P=0.031). VWR increased hepatic glycogen content and increased glucagon-stimulated glycogen depletion, regardless of age (P<0.01). Results from these studies suggest that exercise training can enhance liver glucagon action in aging mice without otherwise altering glucose homeostasis.

**New and Noteworthy:** Few studies have examined the impact of aging on glucagon sensitivity. Here we show that glucagon sensitivity declines from young adulthood to middle age. Yet, 12 weeks of voluntary wheel running, an exercise intervention without the stress of forced treadmill running or swimming, restores glucagon sensitivity in middle-aged male mice without otherwise altering glucose homeostasis.

## 1. Introduction

Aging is associated with an increase in the incidence of metabolic disease, such as type 2 diabetes, obesity, and cardiovascular disease (1-3). Insulin resistance, a key driver of metabolic disease, can be ameliorated through exercise training in both humans (4, 5) and laboratory rodents (6, 7). The counterregulatory hormone to insulin, glucagon, is equally critical in maintaining both glucose and lipid homeostasis, particularly during exercise. While there are limited reports identifying the role of aging in hepatic glucagon sensitivity, those that have studied the intersection between age and glucagon physiology suggest an age-related decline in hepatic glucagon sensitivity (8, 9). Exercise training can improve hepatic glucagon sensitivity in aging rats (9) and in young, healthy male humans (10). Exercise is also a glucagon sensitizing intervention in younger animals (9, 11-15). The bulk of studies demonstrating the glucagon sensitizing effects of exercise in rodents have employed rigorous and carefully dosed exercise protocols, such as treadmill running or swimming. Voluntary wheel running (VWR) in the mouse closely mimics voluntary daily exercise in humans (16), eliminates the stress associated with forced treadmill or swimming exercise interventions (17), better mimics the natural running pattern of a mouse (18) and allows animals to exercise to match their normal circadian cycle (18).

Studies investigating the impact of VWR on improvements in metabolic function in rodents commonly examine the potential to combat the metabolic disturbances caused by diet-induced obesity. In obese mice, exercise decreases adiposity and (19) improves glucose clearance and insulin sensitivity (6, 20, 21).

Aging is associated with increased adiposity and worsened glucose homeostasis (1-3). We set out to assess the effects of aging from young adulthood to middle age on glucagon sensitivity. We further assessed the effects of 12 weeks of voluntary wheel running on glucagon sensitivity when initiated at young adulthood (6 months of age) and middle age (12 months of age).

## 2. Materials and Methods

### Animal Studies

All animal procedures were approved by the Institutional Animal Care and Use Committee of the University of Arizona (IACUC protocol 18-478). All experimental procedures were performed according to the National Institutes of Health Guide for the Care and Use of Laboratory Animals. Male C57BL/6NCrl mice were obtained from the National Institute on Aging mouse colony (Charles River, Wilmington, MA) 1 month prior to the initiation of studies. Mice were maintained on a 14-hour light/10-hour dark cycle in the University of Arizona Health Sciences temperature (22–24°C) and humidity (40– 60%) controlled vivarium. Mice were maintained on a 14-hour light/10-hour dark cycle with unlimited access to food (NIH-31) and water. To acclimate the mice to individual housing, they were singly housed 1 week prior to the beginning of studies and assigned to either remain sedentary or voluntary wheel running (VWR) exercise training.

6- (young), and 12- (middle-aged) month-old mice were either given access to a voluntary running wheel (VWR exercise trained group) or kept in normal housing (sedentary group) for 12 weeks. All mice were housed in the same room, regardless of experimental group for the duration of the study. Body weights (BW) were recorded weekly at 5:00 PM for the duration of the study. Body composition was measured via MRI (EchoMRI™, Houston, TX) at baseline and post exercise training intervention.

Mice provided running wheels were housed in pans outfitted with a freely rotating 12.7cm-diameter wheel with a 5.7cm wide running surface equipped with an electronic counter to measure revolutions (Model 80820F Lafayette Instrument, Lafayette, IN). A motion sensor detected movement of the running wheel in 2 second intervals. The wheel provides virtually friction-free running with < 3 grams of wheel drag. Mice had unrestricted access to the running wheel 24 hours per day for 12 weeks. Distance and time of voluntary running was recorded digitally each day over the 12-week study. Daily running wheel activity data was recorded via AWM monitoring system (Lafayette Instruments, Lafayette, IN). MatLab2021a Software (MathWorks, US) was used to analyze running distance and duration over each 24h period.

### In vivo Measures of Glucose Homeostasis

*In vivo* tests of glucose homeostasis were performed after 10 weeks of VWR. Prior to our *in vivo* metabolic tests, we fasted the mice for 4 hours, 9 AM – 1 PM. To assess glucose clearance, an oral glucose tolerance test (OGTT) was performed. All mice were given an oral glucose gavage (2.5 g/kg BW glucose). Blood was collected via tail vein at baseline and 15 minutes after the oral glucose gavage to assess baseline and oral glucose stimulated insulin (OGSIS), respectively. Blood glucose was read with a glucometer (Bayer Contour Next; Berlin, Germany) at baseline, 15-, 30-, 60-, 90-, and 120-minutes post oral glucose gavage.

An insulin tolerance test (ITT) was performed to assess insulin sensitivity. All mice were given an intraperitoneal (IP) injection of insulin (0.5 IU/ kg BW). Blood was collected at baseline and 15 minutes after the IP insulin injection to assess hypoglycemia stimulated glucagon. Blood glucose was read with a glucometer at baseline, 15-, 30-, 60-, 90-, and 120-minutes post IP insulin injection. The Homeostatic Model Assessment for Insulin Resistance (HOMA-IR) index was calculated using the formula: HOMA-IR=fasting glucose in mmol/l*fasting insulin in μU/ml/22.5 (22).

To assess glucagon sensitivity, a glucagon responsivity test (GRT) was performed. All mice were given an IP injection of glucagon (5 ug/kg/BW; GlucaGen HypoKit, Novo Nordisk, Bagsværd, Denmark). Blood glucose was read with a glucometer at baseline, 15-, 30-, 60-, 90-, and 120-minutes post IP glucagon injection.

### Tissue Collection

Upon completion of the 12-week study, mice were fasted at 9 AM for 4 hours. At 1pm, mice were either given an IP injection of saline or glucagon (20 ug/kg/BW; GlucaGen HypoKit, Novo Nordisk, Bagsværd, Denmark). Mice were sacrificed 15 minutes after injection by decapitation after bell jar isoflurane exposure. Trunk blood was collected into a microcentrifuge tube and clotted on ice for 30 minutes prior to centrifugation at 3,000 x g for 30 minutes. Serum was separated and stored at -80°C. Liver and skeletal muscle (soleus and gastrocnemius combined) were collected, immediately snap-frozen on dry ice, and stored at -80°C until analyses were performed.

### Serum Analyses

Serum glucagon (ELISA; Cat. # 10-1281-01, Mercodia, Uppsala, Sweden) and serum insulin (Cat. # 80-INSMSU-E10, Alpco) were analyzed by commercially available ELISA kits. Serum glucose (Cat #. G7521, Pointe Scientific Inc., Canton, MI, USA) was analyzed using a colorimetric assay.

### Liver Tissue Analysis

We prepared the liver for analyses by powdering using a liquid nitrogen cooled mortar and pestle. This ensured a homogenous sample. For lipid extractions, as we have previously described (23, 24) 10-15 mg of powdered liver samples were weighed and immediately sonicated in 100uL PBS. 1 mL of 100% Ethanol was added and vortexed for 10 minutes. Samples were then centrifuged at 16,000xg for 5 minutes and supernatant was collected for assay analysis. Hepatic triglyceride content (Cat #. T7532, Pointe Scientific Inc., Canton, MI, USA) was analyzed by colorimetric assay and expressed per gram of initial tissue weighed.

Liver glycogen was assayed according to the protocol of Lo et al (1970) (25) and as we have previously described (26, 27). Briefly, frozen powdered liver was weighed immediately prior to boiling and shaking in 30% KOH saturated with NA_2_SO_4_ for 30 minutes. Glycogen was then precipitated with 95% ethanol, followed by centrifugation at 3000 x g for 30 minutes. Glycogen pellets were isolated from supernatant and dissolved in distilled H_2_O, followed by the addition of 5% phenol. H_2_SO_4_ was then pipetted forcefully into the sample. Samples were incubated 30°C for 20 minutes, transferred to a 96 well plate, and read at 490 nm.

Liver total and phosphorylated glycogen synthase was analyzed via western blot. To prepare lysates, 40-50 mg of powdered liver tissue was homogenized in RIPA buffer (Sant Cruz Biotechnology) with protease and phosphatase inhibitors (Halt inhibitor cocktail Cat#1861280, Thermo Scientific). Total protein was quantified by BCA protein assay (Thermo Scientific). Forty micrograms of protein per sample were loaded on a 4-12% Bis Tris gel (Invitrogen) and transferred onto a nitrocellulose membrane (0.2 um pore size). Membranes were blocked for 1 hour at room temperature in a blocking solution (Tris-buffer saline with 0.5% Tween 20 (TBST) with 5% nonfat dry milk) while on rotating shaker at 13 rpm. Membranes were washed 3 times for 5 minutes each with TBST, then incubated overnight at 4°C with primary antibodies diluted 1:1000 in blocking solution (Phospho-Glycogen Synthase (Ser641) Antibody, Cell Signaling cat #3891 and total Glycogen Synthase Cell Signaling Cat #3886). The following day, membranes were washed 3 times for 5 minutes with TBST, then incubated with a secondary antibody (Anti-Rabbit IgG, HRP linked antibody, Cell Signaling #7074S) diluted to 1:2000 in blocking solution for 1 hour at room temperature on rotating plate shaker at 13 rpm. Signals were visualized using SuperSignal West Femto Maximum Sensitivity Substrate (Thermo Scientific) on the Azure Imaging Systems (Azure Biosystems Version 1.5.0.0518) and band intensities were quantified using AzureSpot Version 2.2.167.

We extracted RNA using TRIzol™ Reagent. (Thermo Fisher Scientific, Waltham, MA) following manufacturer protocol and as we have previously described (23, 24). To eliminate phenol contamination, immediately following extraction, RNA was washed with water-saturated butanol and ether (3 times each) using the method of Krebs et al. (2009) (28). We performed cDNA synthesis and RT-qPCR to assess mRNA expression of glycogen synthase as previously described (23, 24) using beta-actin (Actb) as a housekeeping gene. Primer sequences used and annealing temperatures are listed in table 1.

**Table 1.**
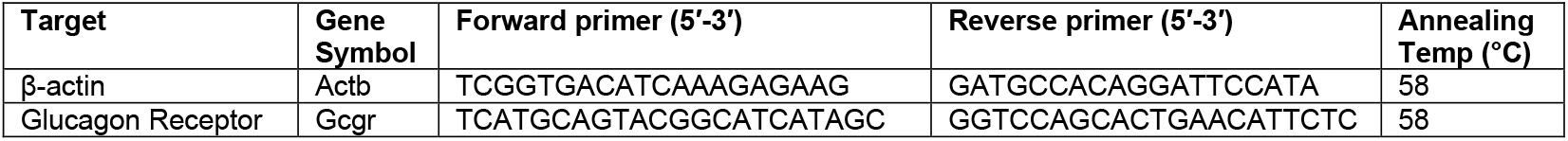
List of Mus musculus Primer Sequences for RT-qPCR.

### Statistical Analyses

Statistical analyses were performed in GraphPad Prism Version 10.4.2 (GraphPad Software). To assess the effect of exercise training within age group (young and aged) on all dependent variables in our animal studies, we used the mixed model procedure. When statistically significant interactions were found, Tukey’s adjustment for multiple comparisons was used to identify significant mean differences. For studies examining the effects of age or exercise only within group, independent t-tests were used. Raw data were plotted in GraphPad Prism Version 10.4.2. All data are presented as mean ± SEM.

## 3. Results

### The effect of Initiating voluntary wheel running at young or middle age on glucose homeostasis and body composition

We first assessed the effect of age at onset of VWR and found no difference in total daily running volume between 6- and 12-month-old mice (Figure 1A-B). 12 weeks of VWR did not affect body weight when initiated at 6- or 12-months of age (Figure 1C-E). VWR decreased fast mass only when initiated at 6-months of age (Figure 1F). Because exercise training can improve glucose homeostasis in both mice and humans, we next assessed the impact of VWR on oral glucose clearance and insulin tolerance. VWR had no effect on oral glucose clearance when initiated at 6-months (Figures 2A&E) or 12-months of age (Figures 2B&F). Similarly, VWR did not affect basal (Figure 2I) or glucose-stimulated (Figure 2J) insulin, regardless of age. While VWR did not affect insulin sensitivity, as assessed by an insulin tolerance test, we did observe a moderate, though not statistically significant (P=0.06) increase in the area under the curve for insulin tolerance in middle-aged mice. HOMA-IR, an estimate of insulin resistance, was also unaffected by exercise, regardless of age (Figure 2N). Because glucagon secretion increases in response to hypoglycemia, we assessed circulating glucagon prior to insulin injection (basal) and 15 minutes post insulin administration (hypoglycemia-stimulated). VWR did not affect basal (Figure 2K) or hypoglycemia stimulated (Figure 2L) glucagon, regardless of age. In line with these findings, the glucagon to insulin ratio was unaffected by VWR in both young and middle-aged mice (Figure 2M).

**Figure 1:**
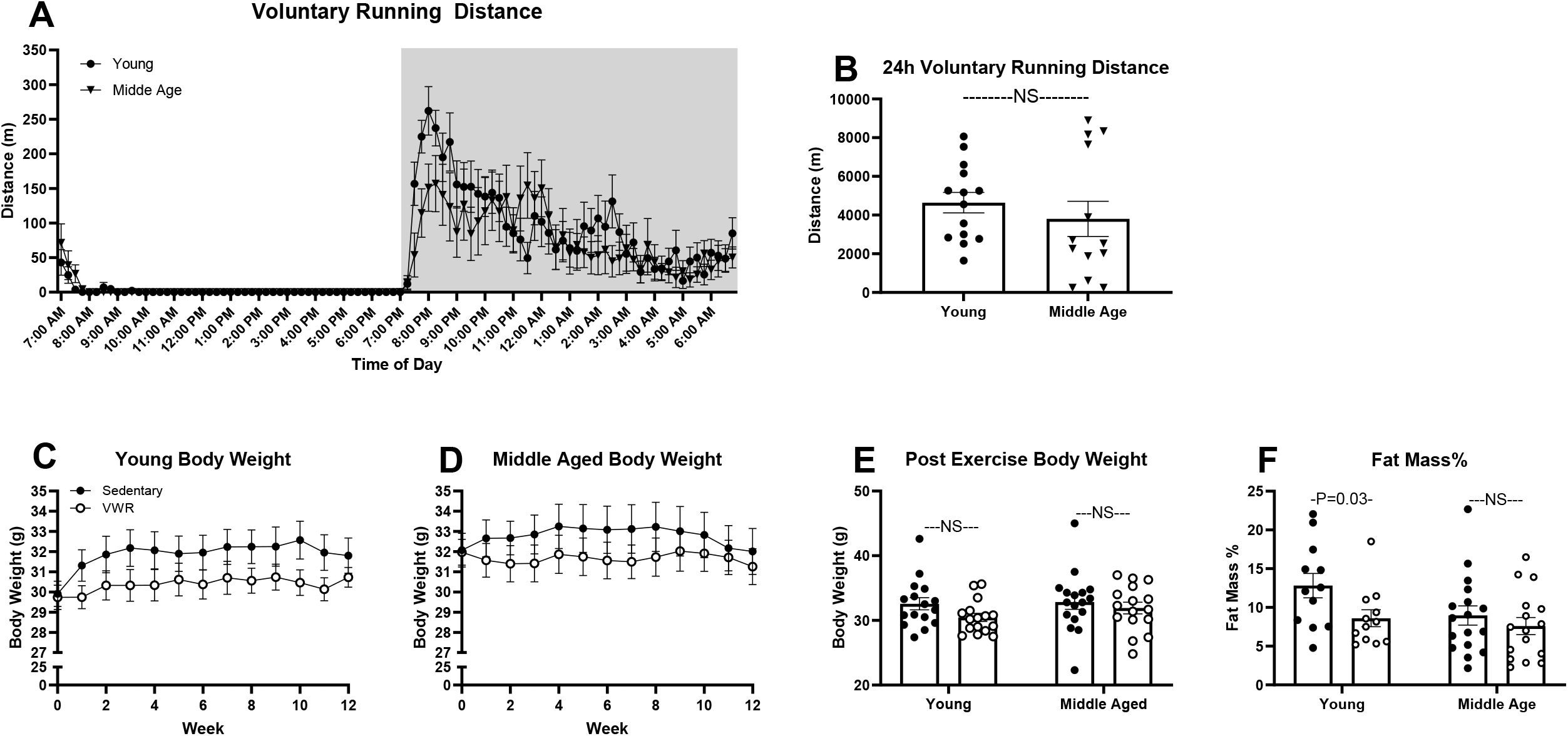
Running volume, body weight, and fat mass over 12 weeks of VWR. Daily running distance in 15-minute intervals (A) and 24-hour total distance ran (B) on week 10 of the exercise training period. Body weight over the 12-week running period (C&D) and final body weight (E) and fat mass percentage (F) at week 10 in young and middle-aged sedentary versus VWR mice. Young sedentary (n=11-16), young exercise trained (n=9-16), middle-aged sedentary (n=12-17), middle-aged exercise trained (n=10-16). Total running distance (B) analyzed with unpaired t-test. Not significant (NS); data are means ± SEM. Voluntary wheel running (VWR).

**Figure 2:**
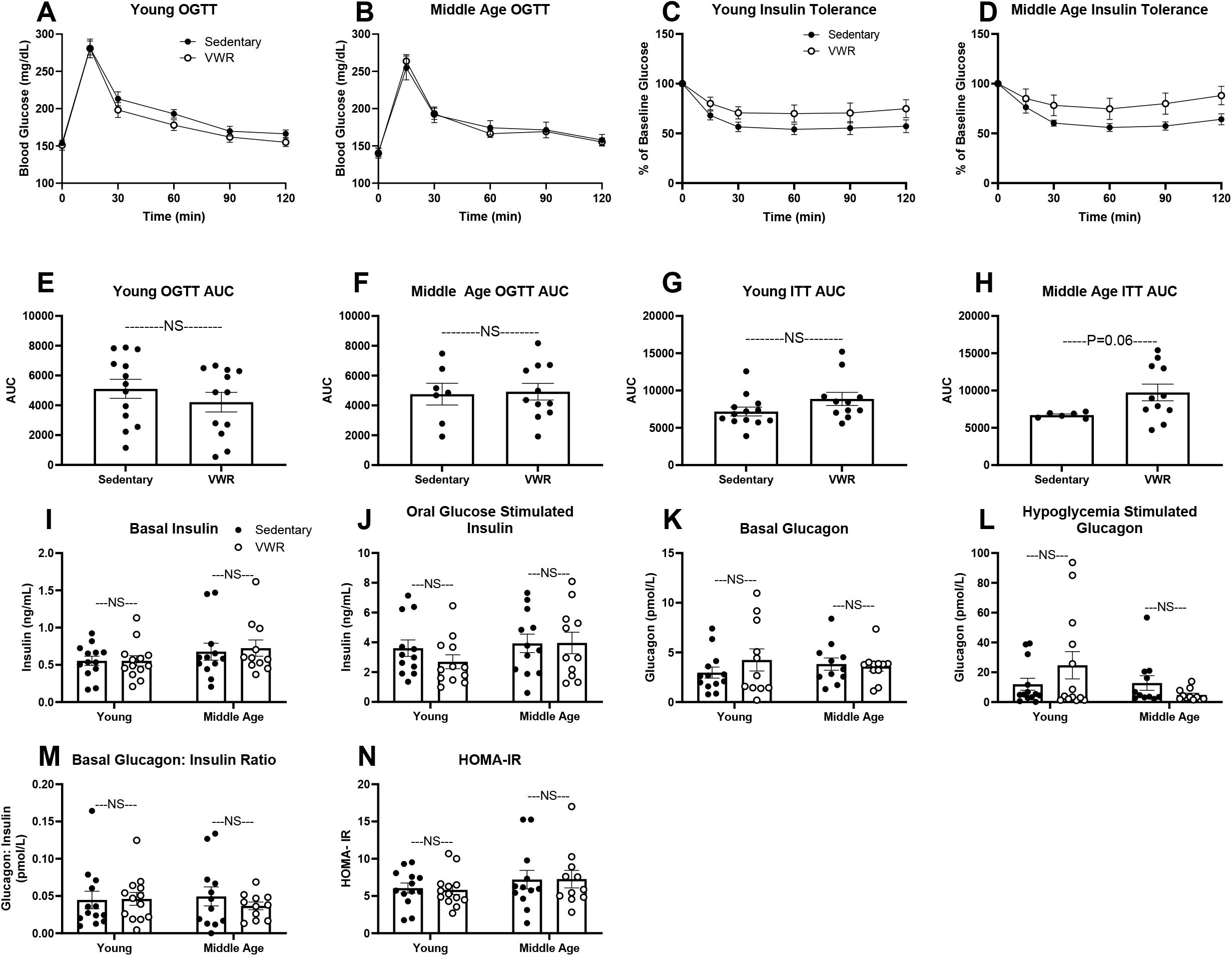
Glucose homeostasis after 10 weeks of exercise training via VWR. Oral glucose clearance (2.5g/kg BW glucose) and corresponding area under the curve in young (A&E) and middle-aged (B&F) sedentary versus VWR mice after 10 weeks of VWR. Insulin tolerance (0.5IU/kg BW) expressed as % change from baseline and corresponding area under the curve in young (C&G) and middle-aged (D&H) sedentary versus VWR mice. Basal (I) and oral glucose-stimulated insulin (J) during OGTT. Basal (K) and hypoglycemia stimulated glucagon (L) during ITT in young and middle-aged sedentary versus VWR mice. The glucagon: insulin ratio (M) and HOMA-IR (N) calculated from basal (4h fasted) glucagon, insulin, and glucose. Young sedentary (n=13), young exercise trained (n=12-13), middle-aged sedentary (n=11), middle-aged exercise trained (n=11), aged sedentary (n=11-13), Differences between sedentary and VWR within age group were analyzed with an unpaired t-test. NS, not significant; data are means ± SEM. Voluntary wheel running (VWR).

### The effect of voluntary wheel running on hepatic lipid accumulation and glycogen content

Excess liver fat impairs glucagon sensitivity at the liver in both humans and rodents (29, 30), thus we next examined the impact of VWR on hepatic lipid accumulation. In line with our observation that VWR only decreased whole-body fat percentage when initiated at 6 months of age, VWR decreased liver triglyceride only when initiated at 6 months of age (Figure 3A). Liver glycogen content increased in response to VWR, regardless of age (Figure 3B). However, skeletal muscle glycogen content only increased in response to VWR when initiated at 6 months of age (Figure 3C).

**Figure 3:**
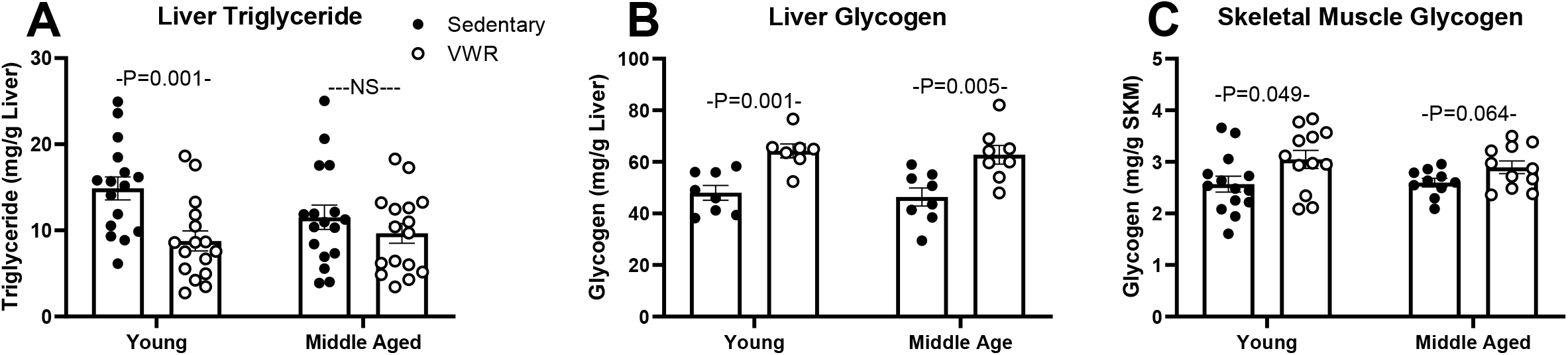
Effect of aging and exercise training on liver triglyceride (A), liver glycogen (B), and skeletal muscle glycogen (C). Young sedentary (n=8-16), young exercise trained (n=7-16), middle-aged sedentary (n=8-17), middle-aged exercise trained (n=8-16). Differences between sedentary and VWR within age group were analyzed with an unpaired t-test. NS, not significant; data are means ± SEM. Voluntary wheel running (VWR).

### The effect of voluntary wheel running on glucagon sensitivity

To assess the impact of VWR on glucagon sensitivity we next performed a glucagon responsivity test after 10 weeks of VWR. We first examined the effects of age on glucagon stimulated hyperglycemia and found that glucagon sensitivity was decreased in sedentary middle-aged mice compared to that of young adult sedentary mice (P=0.046, Figure 4A&B). While VWR had no effect on glucagon stimulated hyperglycemia when initiated at young adulthood (6 months of age) (Figure 4C&D), VWR initiated at middle age (12 months) improved this measure of glucagon sensitivity (P=0.031, Figure 4G&H). Because the initial hyperglycemic effect of glucagon is dependent on glycogenolysis (31) and liver glycogen stores serve as a limited source of glucose, we next assessed liver glycogen content in response to an acute glucagon bolus given intraperitoneally (20 ug/kg/BW). Fifteen minutes after glucagon administration, glucagon decreased liver glycogen content in VWR mice, regardless of the age at which it was initiated. In contrast, exogenous glucagon had no effect on either young adult or middle-aged sedentary mice (Figures 4 E&I). These exercise-induced improvements in hepatic glucagon sensitivity are independent of changes in the mRNA expression of the glucagon receptor (Gcgr), as VWR had no effect on Gcgr mRNA expression at the liver, regardless of age. (Figure 4F&J).

**Figure 4:**
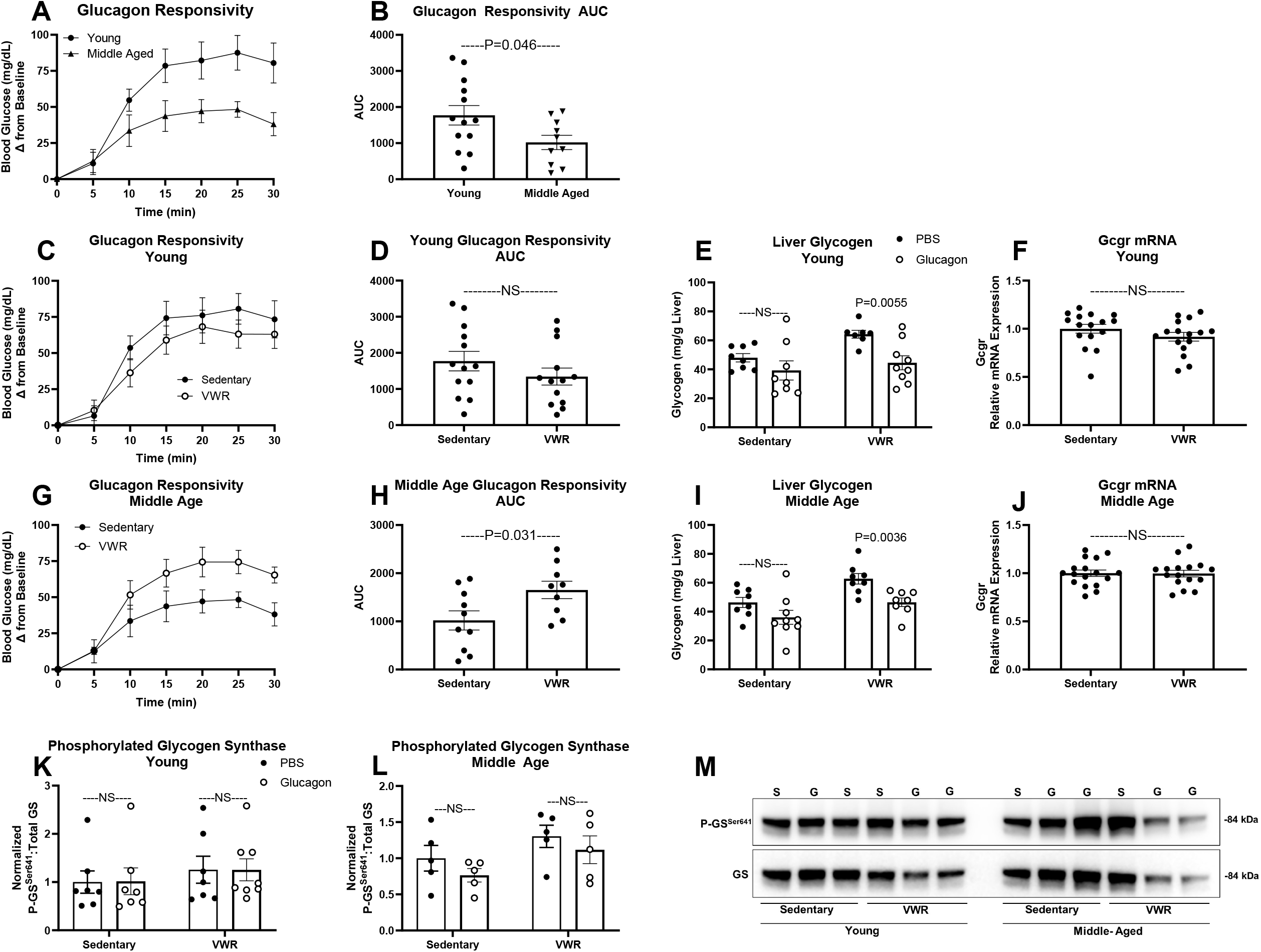
Glucagon Responsivity. Effect of aging and exercise training on glucagon stimulated hyperglycemia in young and middle-aged sedentary male mice (A&B), glucagon stimulated hyperglycemia in young (C&D) and middle-aged (G&H) sedentary versus VWR exercise trained mice. Glucagon-stimulated liver glycogen depletion in young (E) and middle-aged (I) sedentary and exercise trained mice. Glucagon receptor mRNA expression in young (F) and middle-aged (J) sedentary and exercise trained mice. Protein expression of phosphorylated (P-GS^Ser641^) to total glycogen synthase (GS) in young (K) and middle-aged (L) sedentary and exercise trained mice, representative blot (M). Differences between sedentary and VWR within age group were analyzed with an unpaired t-test (A-D and G-H). Differences between PBS and glucagon administration were analyzed with an unpaired t-test (E-F, I-J, and K-M). Not significant (NS); data are means ± SEM. Voluntary wheel running (VWR).

Glucagon signaling at the liver inhibits glycogen synthesis and promotes glycogenolysis through PKA mediated phosphorylation of glycogen synthase and glycogen phosphorylase (32). We quantified phosphorylated and total glycogen synthase as a possible indicator of glucagon action at the liver. We did not detect an effect of either VWR or exogenous glucagon administration on the phosphorylation of glycogen synthase, regardless of age at which exercise was initiated (Figure 4K-M).

## 4. Discussion

Moderate aerobic exercise training is a potent intervention for the treatment and prevention of age-related metabolic disease. Although most research aimed at understanding the metabolic benefits of exercise training in aging humans (4, 5) and mice (6, 7, 19-21) have focused on improvements in insulin sensitivity (4, 5), the role of glucagon signaling is equally critical in the metabolic response to exercise training (31, 33). We have shown that glucagon receptor signaling at the liver plays a critical, protective role in aging (27). Yet, few studies have examined the impact of aging on glucagon sensitivity (8, 9) or how exercise may augment age-related changes in glucagon sensitivity. Herein, we present findings that suggest glucagon sensitivity decreases from young adulthood to middle age and that exercise training in the form of voluntary wheel running (VWR) can ameliorate this age-related decline.

The glucagon sensitizing effects of exercise training in rodents are well documented (9, 11-15), but primarily rely on forced exercise protocols, such as treadmill running and swimming. Because VWR is voluntary and allows mice to maintain a normal diurnal pattern of activity (18), we aimed to assess if VWR similarly enhanced glucagon sensitivity. Similar to that of Bonjorn and colleagues (2002) who reported an increase in glucagon-stimulated hyperglycemia in exercise trained rats with increased liver glycogen stores, we observed that VWR improves glucagon sensitivity in middle-aged mice (11). Because the age of the animals was not reported by Bonjorn et al. (2002), we cannot compared their results to our observed aging effects (11). Podolin and colleagues (2001) investigated age-related changes in hepatic glucagon sensitivity in 4- (young) 12- (middle-aged) and 22- (aged) month old rats. The authors reported a decline in glucagon sensitivity in aged, but not middle-aged animals, as assessed by a decrease in glucagon stimulated adenylate cyclase activity. The authors further reported that exercise training (treadmill running for 60 minutes/ day 5 days per week) restored glucagon sensitivity. Our data in middle-aged mice are contrary to these findings in that we observed a decline in glucagon sensitivity in middle-aged compared to young adult mice, as assessed by *in vivo* glucagon-stimulated hyperglycemia (Figure 4A&B). However, we both showed that aging induced decreases in glucagon sensitivity can be restored by exercise training (Figure 4G&H; (9)). In line with our observation that VWR enhances glucagon sensitivity, we found that acute glucagon administration decreased liver glycogen content only in VWR mice, independent of the age at which exercise was initiated (Figures 4E&I). Importantly, these exercise-induced improvements in glucagon sensitivity are independent of changes in glucose clearance, insulin sensitivity, insulin (basal and glucose stimulated), or glucagon (basal or hypoglycemia stimulated) (Figure 2), as VWR had no effect on these measures of glucose homeostasis.

Surprisingly, the improvement in glucagon sensitivity we observed in middle-aged mice was also independent of changes in liver fat. Excess liver fat impairs glucagon sensitivity at the liver, while decreasing liver fat improves glucagon sensitivity in both humans and rodents (29, 30). In contrast to young mice, middle-aged mice failed to decrease hepatic triglyceride in response to 12 weeks of VWR (Figure 3A). Most studies investigating the impact of hepatic lipid accumulation on glucagon sensitivity are in humans and rodents with obesity or type 2 diabetes (29, 30). Thus, it is possible that the relationship between liver fat and liver glucagon sensitivity is stronger in obese rather than lean animals.

We observed that hepatic glycogen content increased in response to 12 weeks of VWR regardless of age (Figure 3B), confirming previous findings that exercise training increases liver glycogen content in rodents (11, 34). Increasing liver glycogen stores can improve insulin sensitivity in mouse models of monogenic diabetes (35), while decreased liver glycogen leads to hepatic insulin resistance and excess accumulation of liver fat (36). Importantly, Hughey and colleagues (2004) found that AMP-activated protein kinase (AMPK) signaling at the liver is required for an exercise training stimulated increase in liver glycogen (34). Glucagon signaling at the liver activates AMPK (37). In fact, our recent work demonstrates that hepatic AMPK activity is decreased in mice lacking glucagon receptor signaling at the liver (27), providing further evidence that glucagon receptor signaling is critical to the metabolic adaptations induced by exercise training.

The studies described herein do not identify potential mechanisms that could explain the effects of VWR on glucagon sensitivity. While we did not observe exercise or age-related changes in glycogen synthase activity, additional signaling pathways must be examined, including adenylate cyclase activity and the downstream PKA mediated activation of enzymes that regulate both glucose and lipid metabolism, including glycogen phosphorylase. Glucagon stimulated AMPK and downstream targets should also be examined as potential mediators in exercise induced glucagon sensitivity. Despite these limitations, our studies provide strong evidence that glucagon sensitivity decreases from young adulthood to middle age in male mice. Moreover, our findings that 12 weeks of VWR ameliorates this age-related decline in glucagon sensitivity suggest that VWR can enhance glucagon action at the liver in aging mice without otherwise altering glucose homeostasis.

## Authors’ contributions

J.H.S. and T.J.M. conceived and designed the research. T.J.M., A.V., S.G., and K.R.B performed *in vivo* mouse studies, wet lab experiments, corresponding data analysis, and prepared figures. T.F. performed wet lab experiments. T.J.M. and J.H.S. analyzed data, interpreted results of experiments, and prepared figures. T.J.M and J.H.S drafted the manuscript. K.R.B, A.V., S.G., and T.F. edited and revised the manuscript. J.H.S. approved final version of the manuscript.

## Figure Legends

**Supplemental Figure 1:**
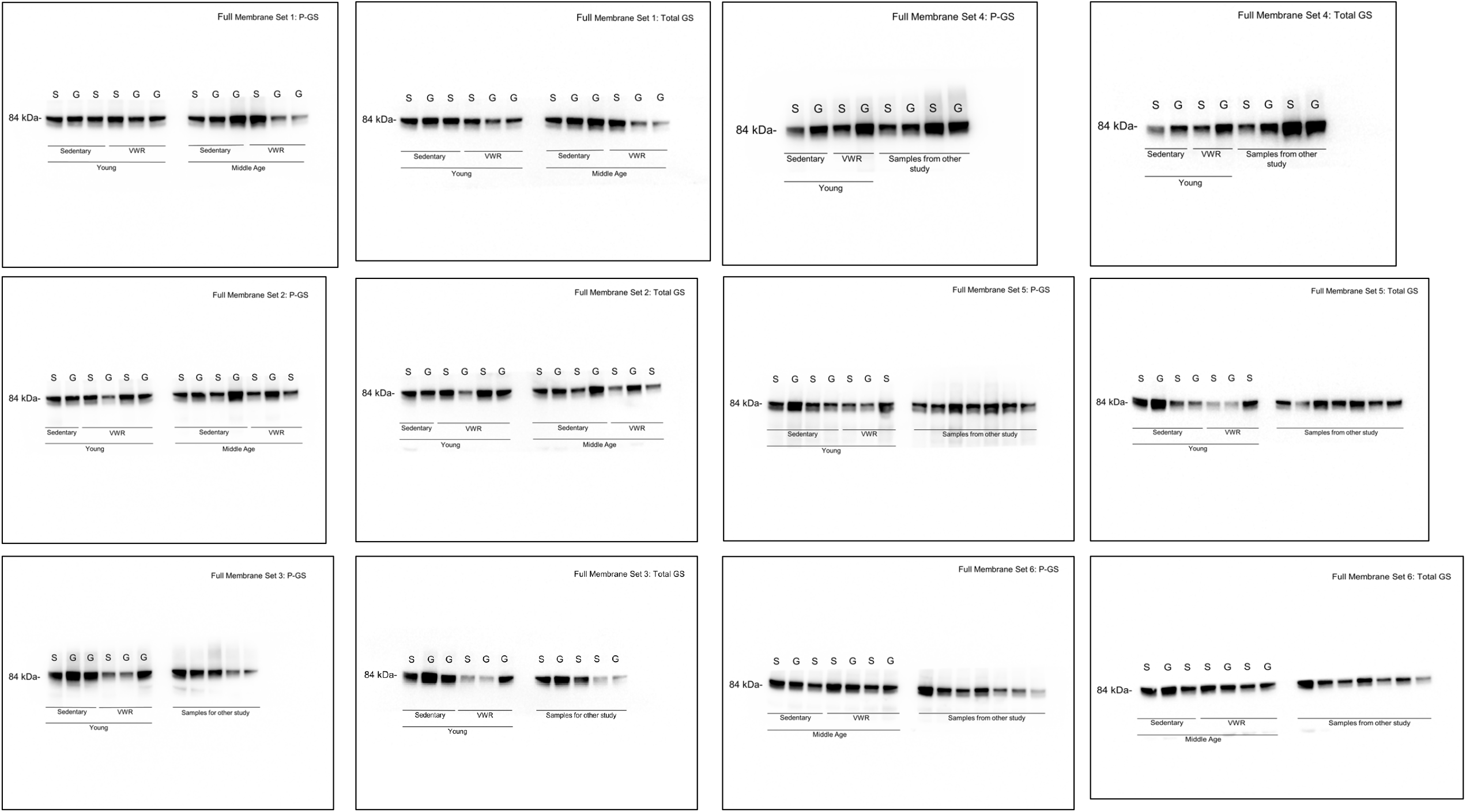
Full Western Blot Membranes from Figure 4M, Phosphorylated and total Glycogen Synthase blots.

